# Identification of novel mutations contributing to azole tolerance of *Aspergillus fumigatu*s through *in vitro* exposure to tebuconazole

**DOI:** 10.1101/2020.07.20.213256

**Authors:** Takahito Toyotome, Kenji Onishi, Mio Sato, Yoko Kusuya, Daisuke Hagiwara, Akira Watanabe, Hiroki Takahashi

## Abstract

Azole resistance of *Aspergillus fumigatus* is a global problem. The major resistant mechanism is a *cyp51A* alteration such as mutation(s) in the gene and the acquisition of a tandem repeat in the promoter. Although other azole tolerances and resistant mechanisms such as *hmg1* mutation are known, few reports describe studies elucidating non-*cyp51A* resistance mechanisms. This study explored genes contributing to azole tolerance in *A. fumigatus* by *in vitro* mutant selection with tebuconazole, an azole fungicide. After three-round selection, we obtained four isolates with low susceptibility to tebuconazole. These isolates also showed low susceptibility to itraconazole and voriconazole. Comparison of the genome sequences of the obtained isolates and the parental strain revealed a non-synonymous mutation in MfsD (Afu1g11820, R337L mutation) in all isolates. Furthermore, non-synonymous mutations in AgcA (Afu7g05220, E535Stop mutation), UbcD (Afu3g06030, T98K mutation), AbcJ (Afu3g12220, G297E mutation), and RttA (Afu7g04740, A83T mutation), a protein responsible for tebuconazole tolerance, were found in at least one isolate. Clarification by constructing the MfsD R337L mutant suggests that the mutation contributes to azole tolerance. Disruption of the *agcA* gene and reconstruction of the A83T point mutation in RttA led to decreased susceptibility to azoles. The reversion of T98K mutation to wild type in UbcD led to the level of azole susceptibility comparable to the parental strain. These results suggest that these mutations contribute to lowered susceptibility to medical azoles and to agricultural azole fungicides.

## Introduction

*Aspergillus fumigatus*, a filamentous fungus commonly found in the natural environment, is known to be the major causative agent of aspergillosis. Its 2–3 μm diameter conidia are dispersed easily in the air, from which they can reach the human lung alveoli. The fungus colonizes and invades lung tissue in immunocompromised hosts or hosts with local immunosuppression in the lung, leading to aspergillosis. Recent global estimates indicate that 3 million cases of chronic pulmonary aspergillosis (CPA) and 0.3 million cases of invasive aspergillosis (IA) occur annually, each with high mortality even if treated (1–3).

Azole antifungals are frontline agents for aspergillosis treatment. Therefore, the emergence of azole resistance in *A. fumigatus* has raised global concern. As demonstrated in earlier studies (4–10), azole-resistant strains often appear during treatment. Resistant-responsible mutations were found mainly in Cyp51A (Erg11), which is the target protein of azole antifungals (11, 12). Actually, *cdr1B* (13), *hapE* (14), and *hmg1* (4) have been known as non-*cyp51A* genes responsible for azole resistance in clinical isolates. Nevertheless, related reports are few.

Azole antifungals have also been used worldwide as agricultural fungicides. The agricultural azole fungicides and medical azoles share the antifungal mechanism.

Tolerance to the fungicides might confer cross-resistance to medical azoles. Consequently, the fungicide residues are thought to act as a selective pressure in the soil, which has led to the appearance of azole-resistant *A. fumigatus* in the environment (15, 16). The prevalent environmental azole-resistant strains also include mutation(s) in *cyp51A* and additionally acquire a tandem repeat (TR) in the promoter (17). Tebuconazole (TBCZ) is a widely used agricultural fungicide. Ren et al. (18) and Cui et al. (19) reported that TBCZ selection induced *cyp51A* mutations with TR in the promoter. Zhang et al. also induced azole resistance *in vitro* through the pressure of fungicides including TBCZ. The mutations induced in the obtained isolates were found through genome sequencing (20). However, few such studies have been conducted to shed light on responsible genes for azole tolerance and resistance.

For this study, we induced mutations with TBCZ selection pressure *in vitro*, demonstrating that four isolates with lowered susceptibility to TBCZ were obtained. These isolates also showed lowered susceptibility to medical azoles as well as TBCZ. Furthermore, three genes contributing to azole tolerance were identified and confirmed.

## Results

### *In vitro* selection of the isolate adapted to TBCZ

We screened *A. fumigatus* isolates using three-round selection for the 1/2 minimum inhibitory concentration (MIC) of TBCZ *in vitro* (Fig. 1A). The conidia of 8, 7, and 4 isolates obtained in each selection round were pooled. Then MIC of TBCZ in the pooled fungi was determined. The pooled conidia were used for the next selection to accumulate additional mutations. The MIC of TBCZ against obtained isolate mixtures increased gradually to 32 μg/mL (Fig. 1B). Finally, four isolates were obtained: TBCZ31, TBCZ33, TBCZ34, and TBCZ37. Also, the MICs of TBCZ were determined (Table 1). The four isolates showed high MIC of TBCZ: 8–32 μg/mL (Table 1).

**Table 1.** Summary of non-synonymous mutation in each gene.

| Selection round | Strains | MfsD<br>R337L | UbcD<br>T98K | AbcJ<br>G297E | AgcA<br>E535Stop | RttA<br>A83T | Afu6g01930<br>G160G<br>(synonymous) | MIC of<br>TBCZ<br>(µg/mL) |
| --- | --- | --- | --- | --- | --- | --- | --- | --- |
| 1 | TBCZ13 |  |  |  |  |  | ND<sup>b</sup> | 4 |
|  | TBCZ14 |  |  |  |  |  | ND<sup>b</sup> | 2 |
|  | TBCZ15 | ○<sup>a</sup> |  |  |  |  | ND<sup>b</sup> | 4 |
|  | TBCZ16 |  |  |  |  |  | ND<sup>b</sup> | 2 |
|  | TBCZ17 |  |  |  |  |  | ND<sup>b</sup> | 4 |
|  | TBCZ18 |  |  |  |  |  | ND<sup>b</sup> | 8 |
|  | TBCZ19 |  |  |  |  |  | ND<sup>b</sup> | 2 |
|  | TBCZ10 |  |  |  |  |  | ND<sup>b</sup> | 4 |
| 2 | TBCZ21 | ○ |  |  |  |  | ND<sup>b</sup> | 16 |
|  | TBCZ23 | ○ | ○<sup>a</sup> |  |  |  |  | 8 |
|  | TBCZ24 | ○ |  | ○ |  |  | ○ | 8 |
|  | TBCZ25 |  |  |  |  |  | ND<sup>b</sup> | 8 |
|  | TBCZ28 |  |  |  |  |  | ND<sup>b</sup> | 16 |
|  | TBCZ29 | ○ |  |  |  |  | ND<sup>b</sup> | 8 |
|  | TBCZ20 |  |  |  |  |  | ND<sup>b</sup> | 8 |
| 3 | TBCZ31 | ○ |  |  | ○ |  |  | 8 |
|  | TBCZ33 | ○ |  |  | ○ |  |  | 16 |
|  | TBCZ34 | ○ | ○ | ○ |  |  | ○ | 32 |
|  | TBCZ37 | ○ |  |  |  | ○ |  | 32 |

Blank cells show no non-synonymous mutation for each gene.
<sup>a</sup> The base called mixed nucleotides mutated and original.
<sup>b</sup> Not determined.

**Figure 1.**
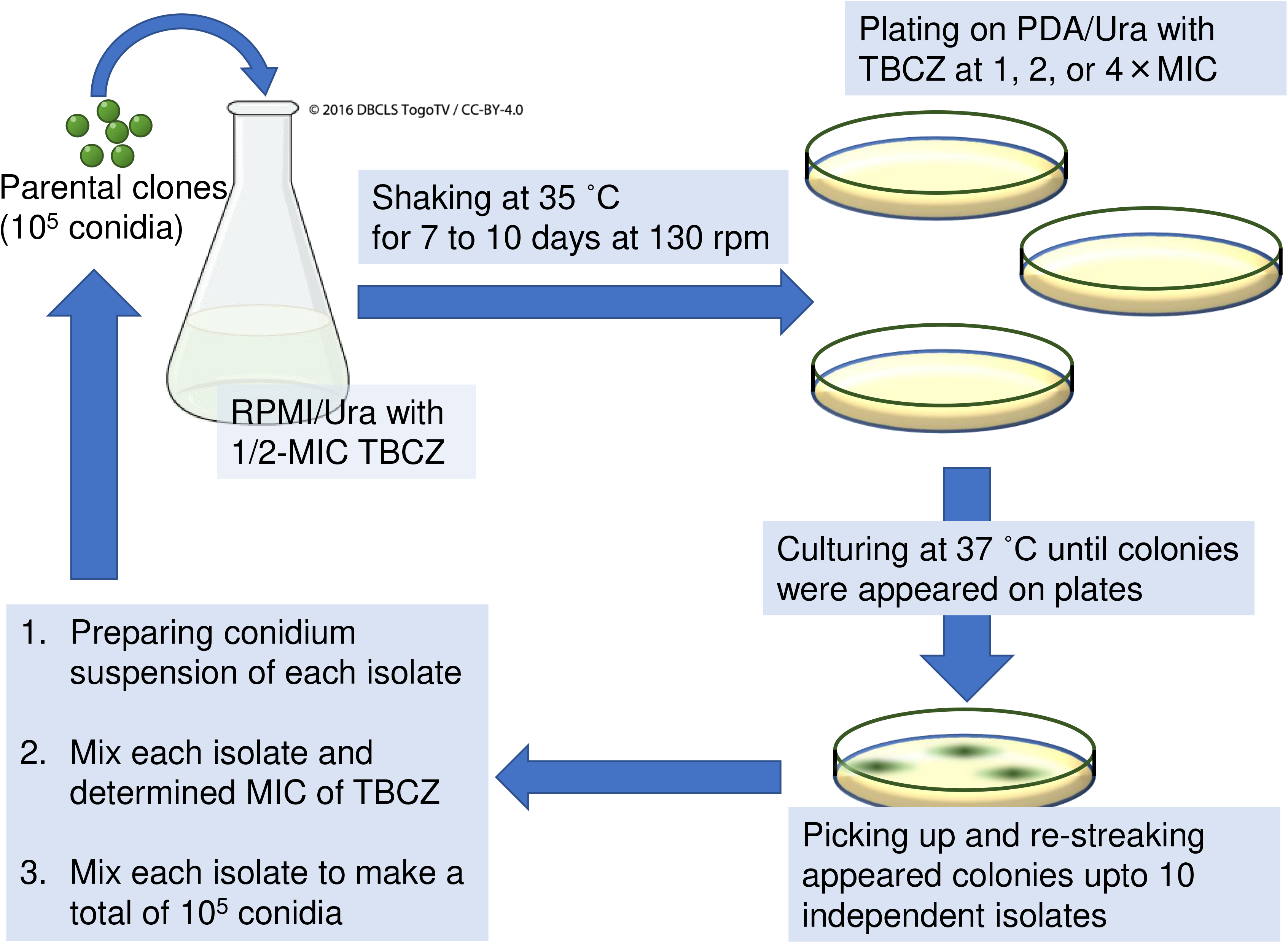
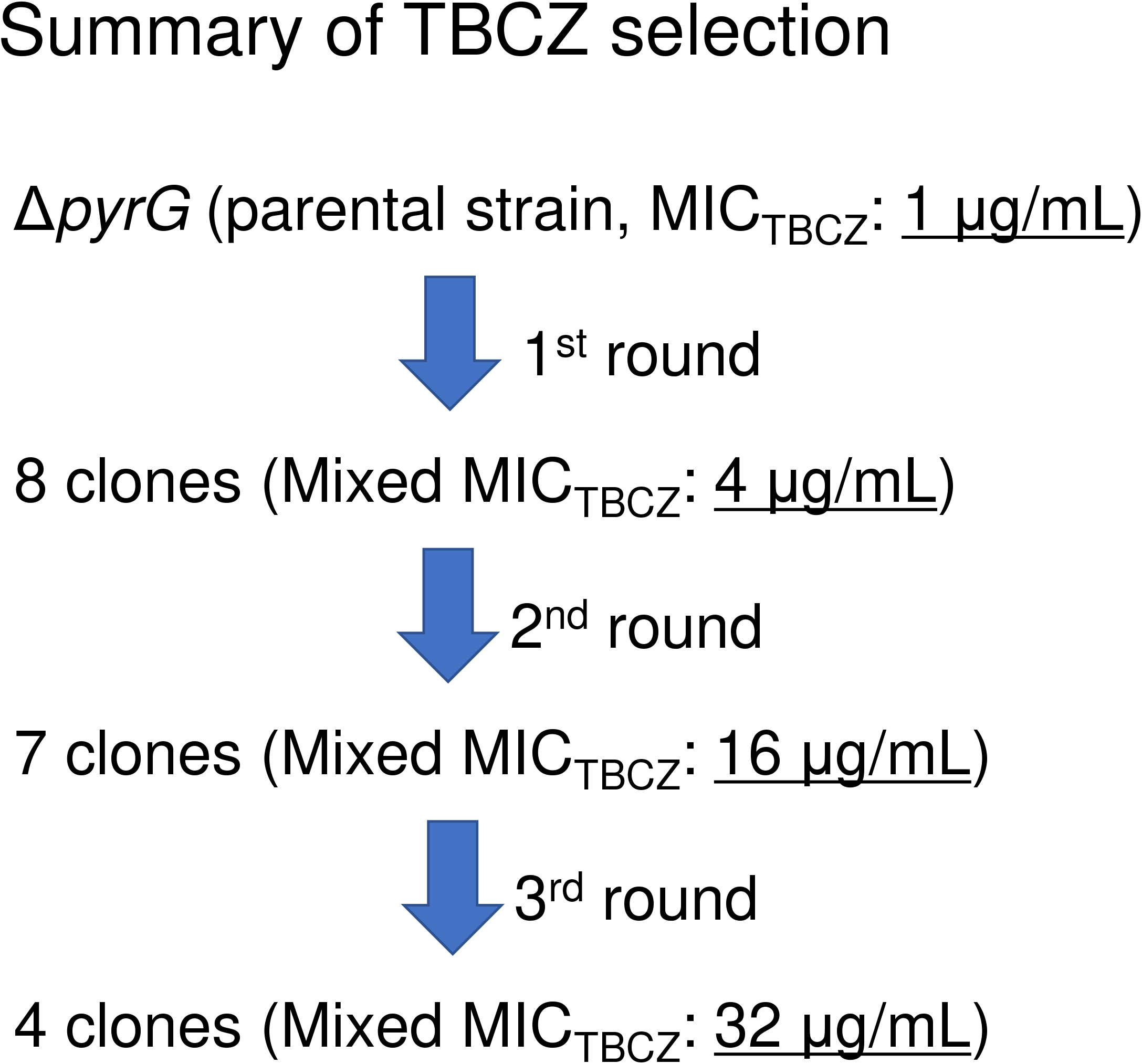
Summary of TBCZ selection. The procedure is presented in *A*. The number of isolates obtained after each selection round and the mixed MICs are shown in *B*.

### Macroscopic phenotype of obtained isolates

The colonies of TBCZ31, TBCZ34, and TBCZ37 are presented in Fig. 2A. The TBCZ34 colony diameter was larger than that of the parental strain (Δ*pyrG*). Also, the TBCZ31 colony diameter was smaller than Δ*pyrG* (Fig. 2). The TBCZ37 growth was comparable to that of the parental strain.

**Figure 2.**
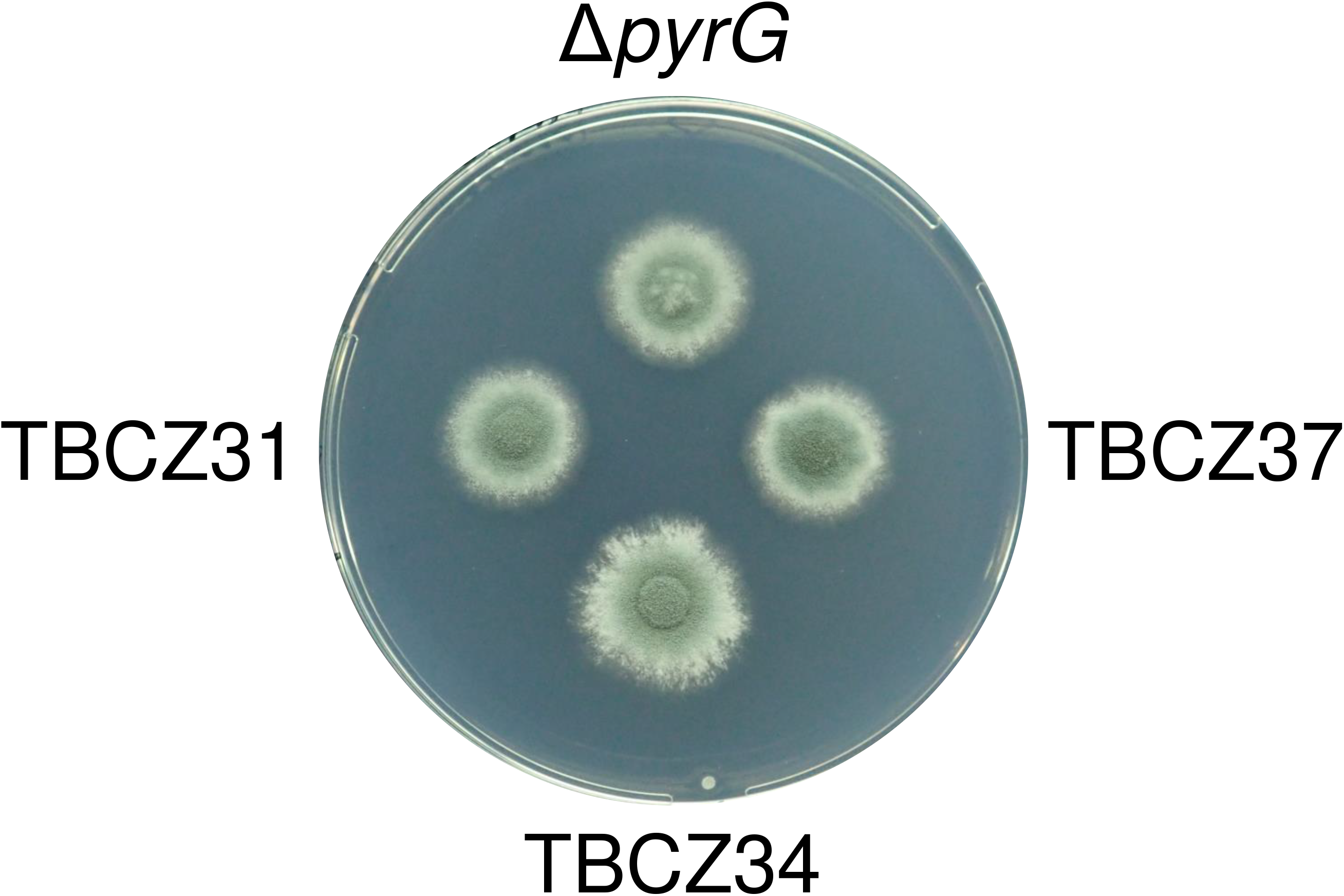
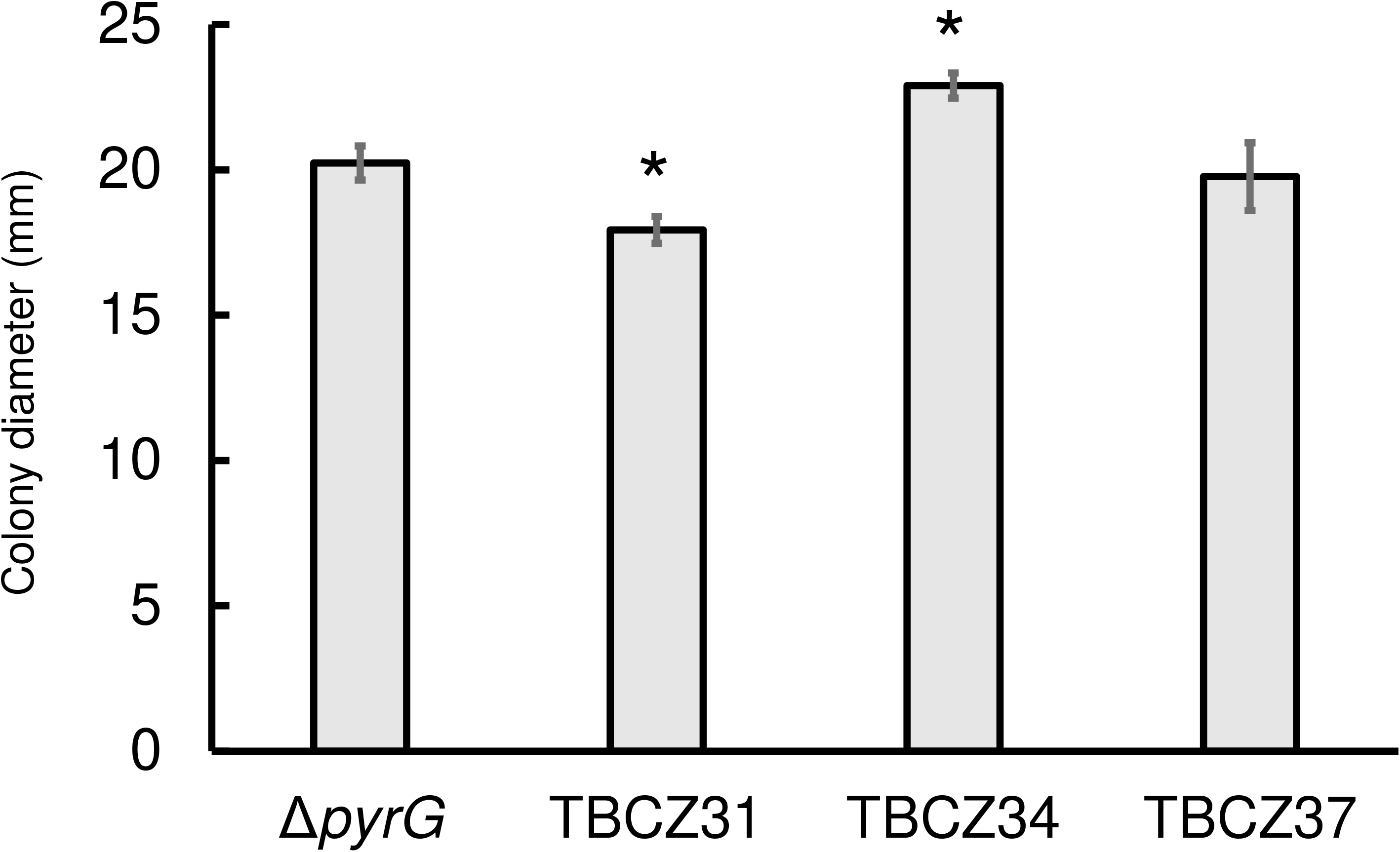
Isolates TBCZ31, 34, and 37 obtained after a three-round selection with TBCZ. After culturing on MM at 35 °C for 96 h, representative macroscopic image and colony diameters are depicted respectively in *A* and *B*. Values in panel *B* are means ± SD of three independent colonies. Asterisks denote significant differences (*p* < 0.05).

### Cross-resistance of TBCZ-adapted isolates to medical azoles

We measured the MICs of medical antifungals, micafungin (MCFG), amphotericin B (AMPH), itraconazole (ITCZ), and voriconazole (VRCZ) against obtained isolates TBCZ31, TBCZ33, TBCZ34, and TBCZ37. Every isolates grew in 4 μg/mL VRCZ (Table 2) and in 2 μg/mL or higher concentration of ITCZ. The MICs of non-azole antifungals MCFG and AMPH were not higher in the isolates (Table 2). These data show that these isolates acquired cross-resistance to the medical azoles via TBCZ selection.

**Table 2.** MICs of antifungals against TBCZ31, TBCZ33, TBCZ34, TBCZ37.

| **Strain** | **ITCZ** | **VRCZ** | **MCFG** | **AMPH** |
| --- | --- | --- | --- | --- |
| *ΔpyrG* | 0.25 | 0.5 | ≤ 0.015 | 1 |
| **TBCZ31** | 2 | 4 | ≤ 0.015 | 1 |
| **TBCZ33** | 2 | 4 | ≤ 0.015 | 1 |
| **TBCZ34** | >8 | 4 | ≤ 0.015 | 1 |
| **TBCZ37** | 2 | 4 | ≤ 0.015 | 1 |
| **TBCZ15** | 0.5 | 2 | ≤ 0.015 | 1 |
| **TBCZ34/*ubcD*<sub>wt</sub>** | 0.25 | 0.5 | ≤ 0.015 | 1 |

**MICs were found at 96 h after inoculation.**

### Non-synonymous mutations are detected in TBCZ-adapted isolates

To identify the mutations affecting azole susceptibility, TBCZ31, TBCZ33, TBCZ34, and TBCZ37 isolates were subjected to genome sequencing. Comparison to the parental strain showed that a non-synonymous mutation in a major facilitator superfamily (MFS) gene *mfsD* (Afu1g11820) was present in all four isolates (Table 1). Non-synonymous mutation in *agcA* (Afu7g05220) was found in TBCZ31 and TBCZ33. Non-synonymous mutations in *ubcD* (Afu3g06030) and *abcJ* (Afu3g12220) were found in TBCZ34. Results of homology analysis show that *ubcD* is a homolog of ubiquitin-conjugating enzyme (E2) gene and that *abcJ* is a member of ATP-binding cassette transporters. Mutations in a hypothetical gene as a responsible gene for TBCZ tolerance *rttA* (Afu7g04740) were identified in TBCZ37. To confirm the step in which each mutation appeared during *in vitro* selection, isolates obtained from the first and the second selection rounds were examined using Sanger sequencing (Table 1). The mutation in *mfsD* was detected in an isolate after the first selection round. Four of seven isolates possessed the mutation after the second round. The T98K mutations in UbcD and G297E in AbcJ were detected respectively in TBCZ23 and TBCZ24 after the second selection round. Mutations in *agcA* and *rttA* were found in no isolate after the second round. These data suggest that R337L mutation in MfsD appeared initially. Then T98K mutation in UbcD and finally the mutation in AgcA or RttA were introduced into the population.

### Growth deceleration by *mfsD* disruption

To elucidate the effects of R337L mutation in MfsD, we investigated the susceptibility of TBCZ15 obtained after the first selection round (Table 1). The growth of TBCZ15 was slower than that of the parental strain (Fig. 3), indicating that the mutation led to slower growth in *A. fumigatus*. As shown in Table 2, the MICs of azoles against TBCZ15 were higher than the parental strain. These results suggest that the mutation in MfsD conferred azole tolerance. Next, we constructed *mfsD* disruptant from *A. fumigatus* AfS35 and examined its susceptibility to antifungals. The disruptants showed marked growth acceleration compared with the parental strain (Fig. 4), also indicating that *mfsD* has some relation with the growth speed, although we were unable to detect any difference between disruptants and the parental strain in terms of susceptibility to azoles (data not shown). These results suggest that the effect of R337L mutation in MfsD differs from that of the absence of the gene.

**Figure 3.**
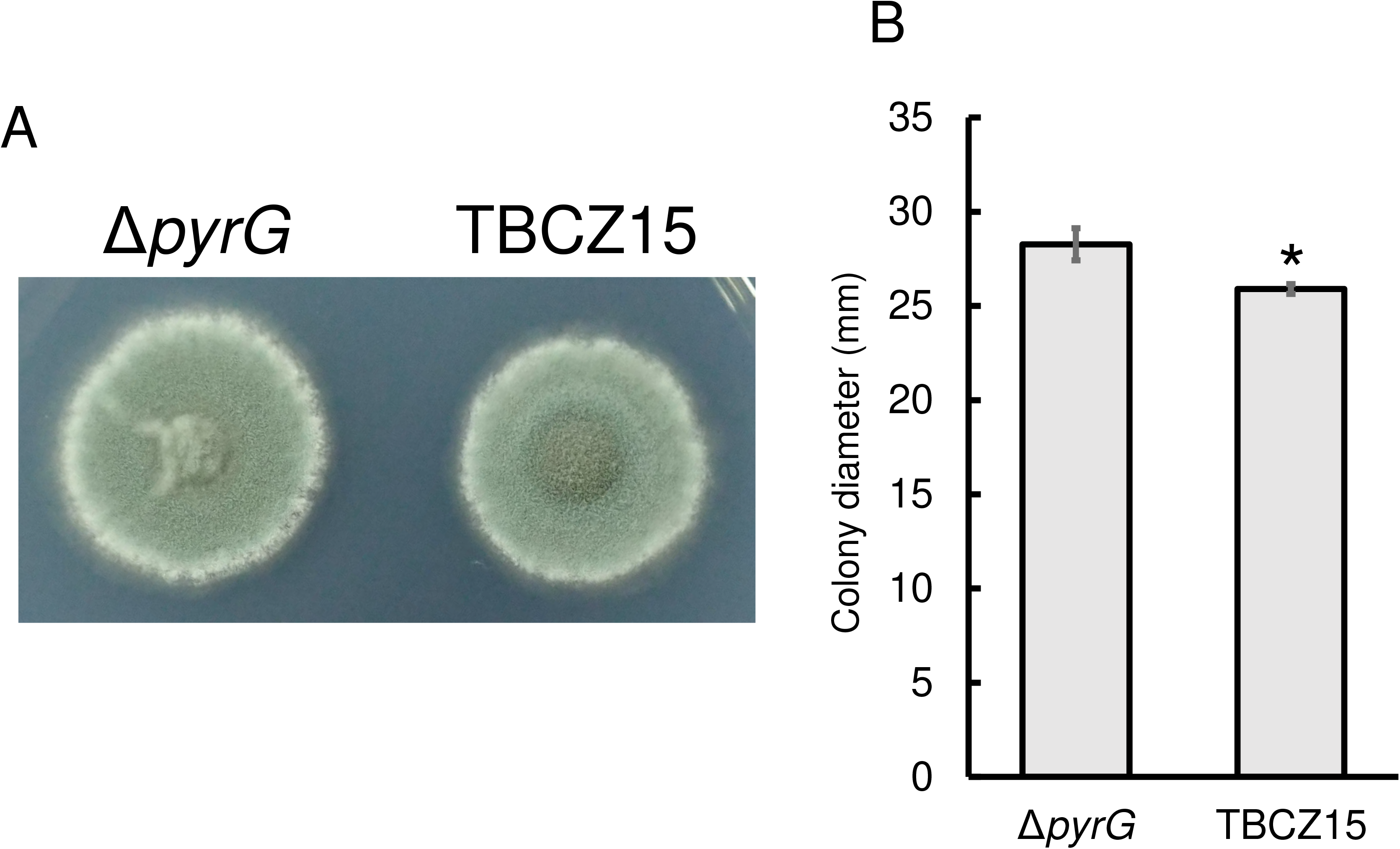
Colonies and diameters of TBCZ15 and the parental strain (Δ*pyrG*) cultured on MM at 35 °C for 120 h. (*A*) The macroscopic image of TBCZ15 and the parental strain. Colony diameters are shown in panel *B*. Values in panel *B* are means ± SD of three independent colonies. Asterisks denote significant differences (*p* < 0.05).

**Figure 4.**
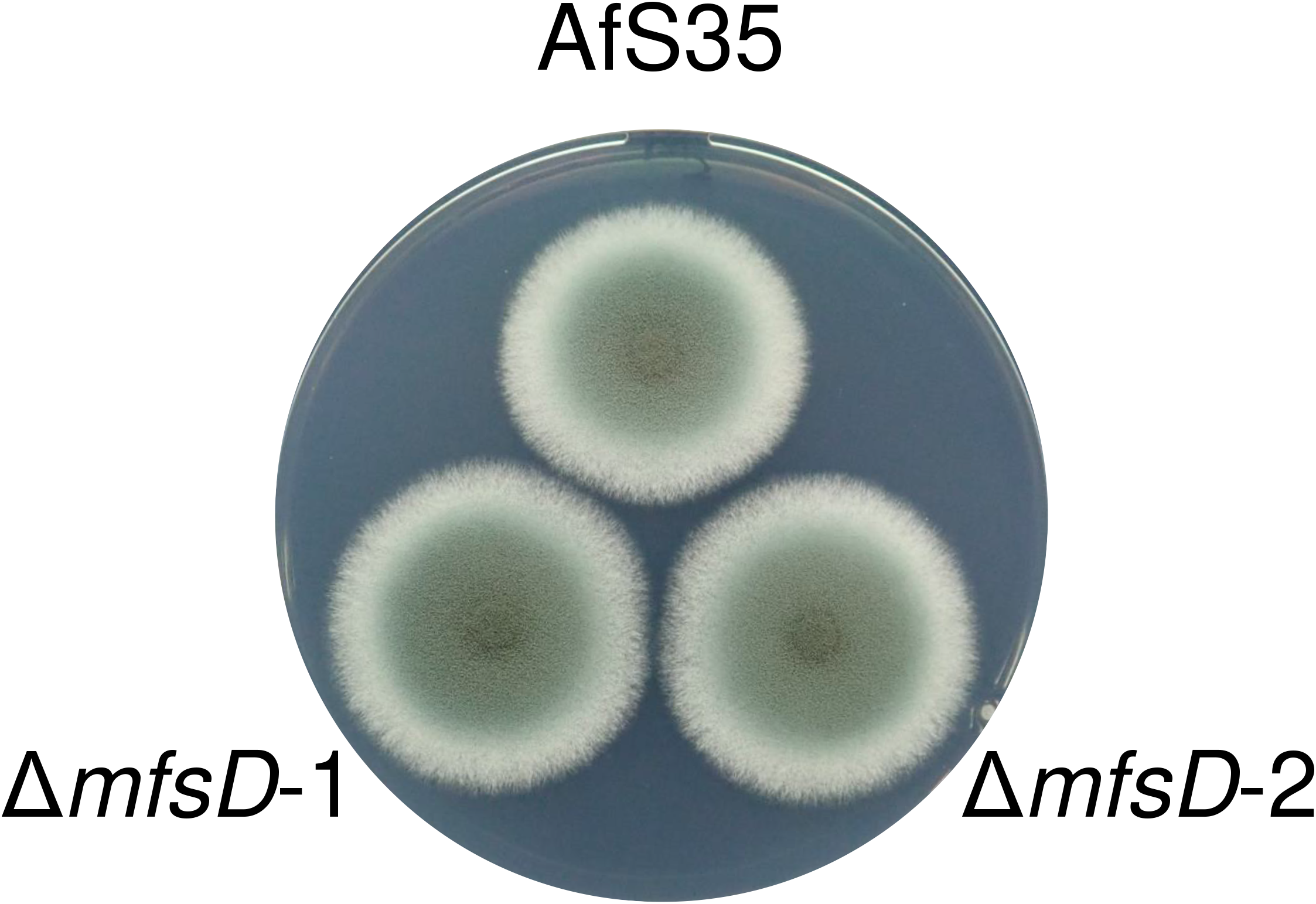
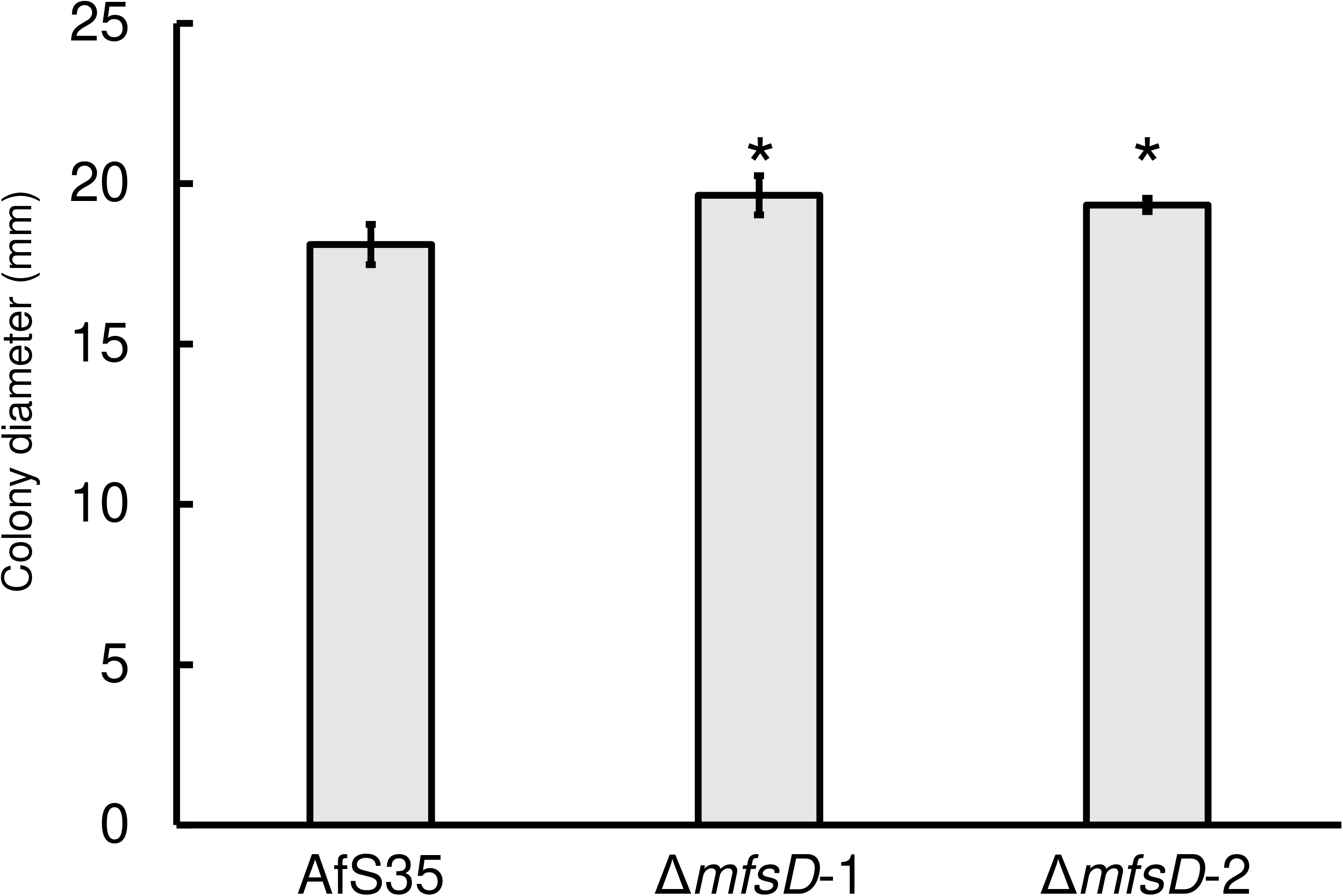
Colony of *mfsD* disruptants. The macroscopic image of two independent *mfsD*-disrupted strains, Δ*mfsD*-1 and Δ*mfsD*-2 and their parental strains AfS35 cultured on MM at 35 °C for 72 h are shown in *A*. Colony diameters measured at 48 h are shown in panel *B*. Values in panel *B* are means ± SD of three independent colonies. Asterisks denote significant differences (*p* < 0.05).

### Reduced susceptibility to azoles in *agcA* disruptant

We then attempted to ascertain whether the other mutation was involved in azole resistance. The mutation in *agcA* found in TBCZ31 and TBCZ33 was a nonsense mutation at the 353rd codon, which caused impairment of 163 amino acids from the C-terminus of AgcA. We hypothesized that the impairment led to functional disruption of AgcA. Therefore, we constructed an *agcA* disruptant by replacing the gene with the hygromycin resistance gene in *A. fumigatus* AfS35. The resulting strain Δ*agcA* exhibits normal growth (Fig. 5). The MIC of VRCZ against Δ*agcA* was 1 μg/mL, which was higher than the MIC against the parental strain (0.5 μg/mL). The result suggests that *agcA* plays a role in azole susceptibility.

**Figure 5.**
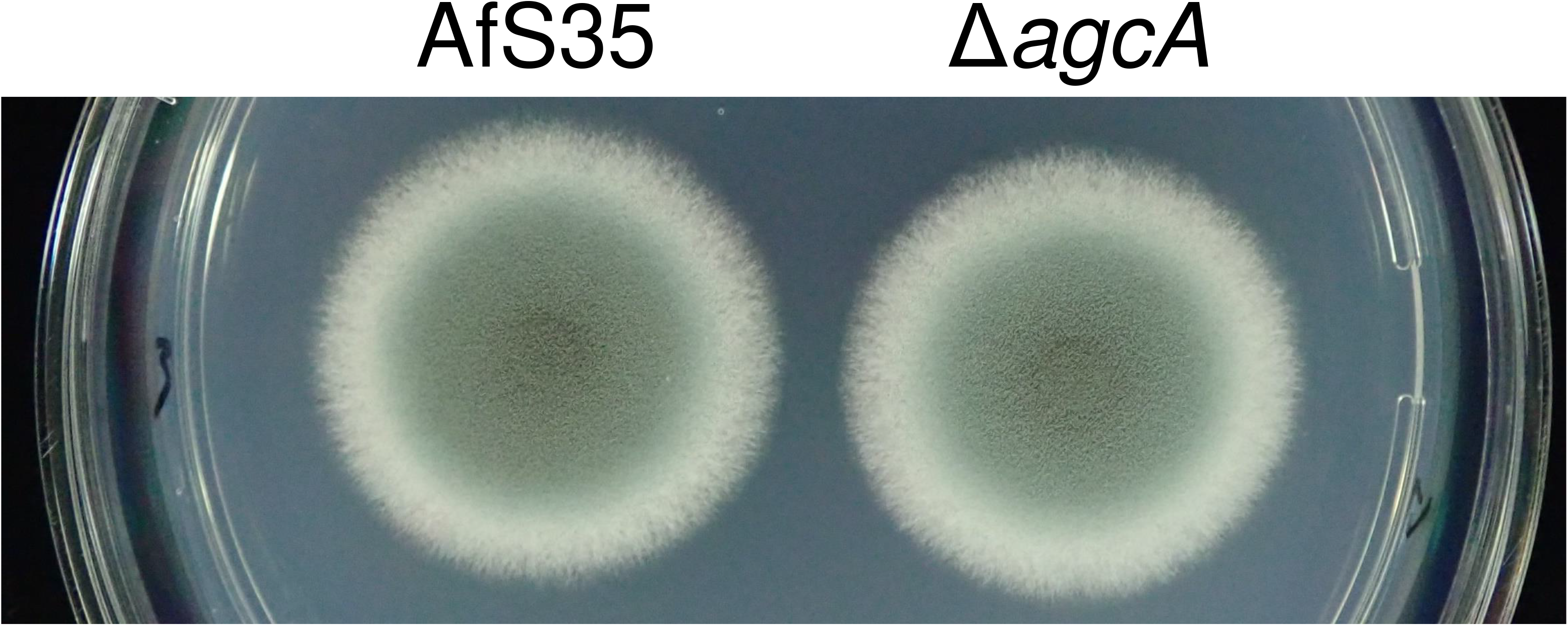
Macroscopic morphology of AfS35 (parental) and agcA disruptant (Δ*agcA*).

### Azole susceptibility reduced by the mutation in *ubcD*

We examined whether the mutation in *ubcD* played a role in azole susceptibility. We tried to disrupt *ubcD* in *A. fumigatus* AfS35, with the result that no transformant was obtained. Instead, we tried to construct a strain for which the mutation of *ubcD* in TBCZ34 was replaced with a wild type *ubcD* gene. The susceptibilities to ITCZ and VRCZ of the resultant isolates were comparable to that of the parental strain (Table 2). This result suggests that the mutation in *ubcD* was responsible for the azole resistance.

### Azole susceptibility reduced by introduction of the mutation in *rttA*

We tried to ascertain whether the mutation in *rttA* plays a role in azole susceptibility. However, we were unable to disrupt *rttA* in *A. fumigatus* AfS35. Therefore, we sought to introduce a mutation (A83T) in RttA of the AfS35 strain (Fig. S4 for the construction). The strain with A83T mutation in RttA (*rttA*_A83T_) was more tolerant of voriconazole (MIC of VRCZ, 0.5 μg/mL) than the control strain (*rttA*_wt_, MIC of VRCZ, 0.25 μg/mL) was, thereby indicating that the mutation contributes to azole tolerance.

## Discussion

In this study, we identified four genes related to azole tolerance: *mfsD*, *ubcD*, *agcA*, and *rttA* genes. For three of those genes (*mfsD*, *ubcD,* and *rttA* genes), the relation with azole tolerance has never been reported. The mutations and the order in which they occurred are shown in Fig. 6.

**Figure 6.**
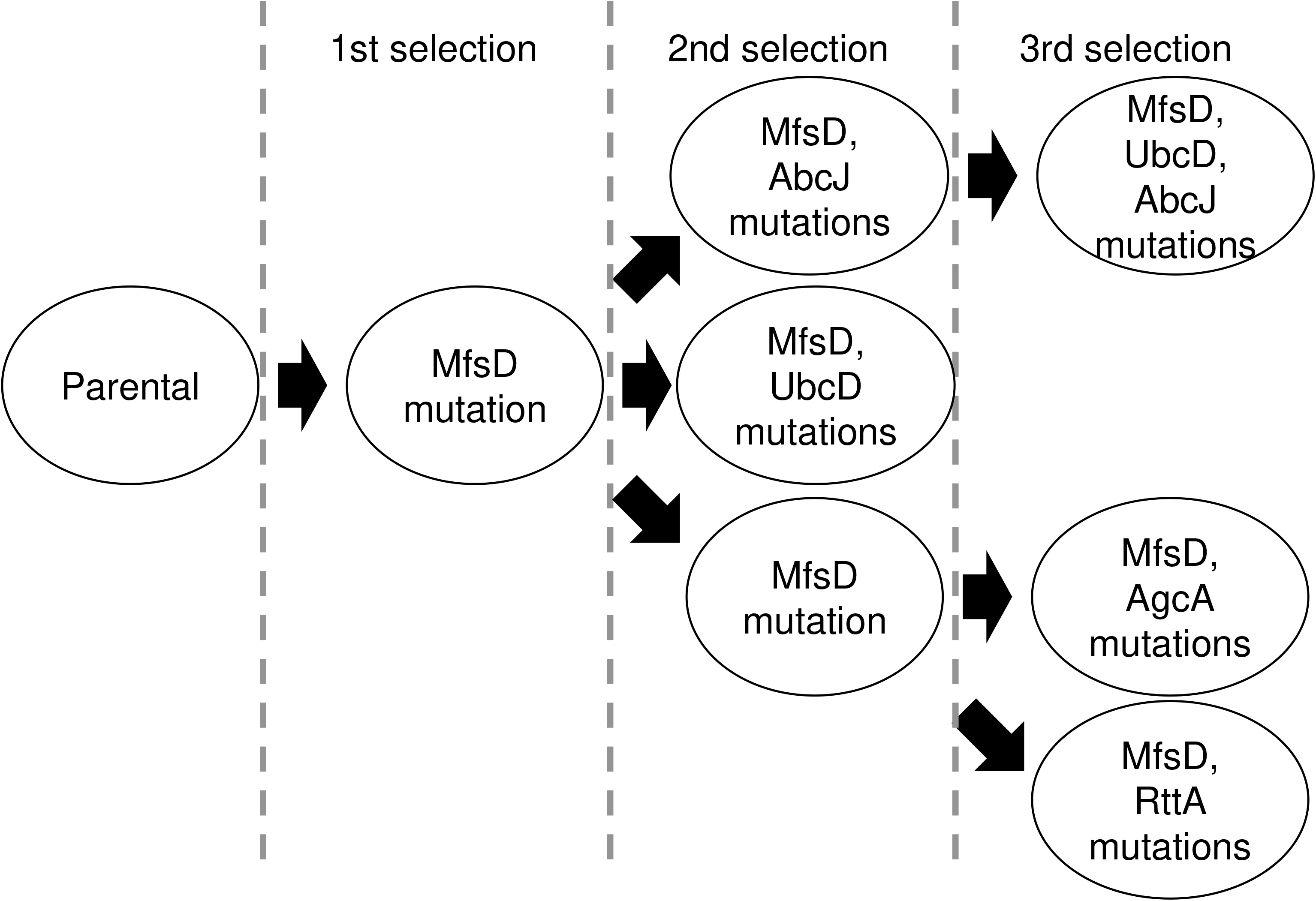
Summary of TBCZ adaptation of isolates obtained in this study.

Here, we selected fungal cells in a liquid medium and then plated the resultant fungal cells on agar plates with various TBCZ concentrations. During selection in the liquid medium, fungal cells formed ball-like structures, suggesting that the balls contained fungal cells of both types before and after mutation, as reported from an earlier study (15).

During the three selection rounds, one strain with R337L mutation in the *mfsD* gene was isolated among eight strains from the first selection round. The mutation was also detected in four of eight strains isolated from the second selection round and all four strains from the third selection round (Table 1). This result indicates that the R337L mutation in MfsD was enriched among the strains. Further analysis is necessary to elucidate the mechanism of how the mutation confers azole tolerance.

The nonsense mutation at the 535th residue in AgcA and disruption of *agcA* caused the azole tolerance phenotype. AgcA homologs are found broadly, for example, in *Saccharomyces cerevisiae* (Agc1p) and human (SLC25A12, also known as Aralar, and SLC25A13, also known as ARALAR2, CITRIN, or CTLN2). These proteins are mitochondrial inner membrane aspartate/glutamate transporters (21, 22) composing the malate–aspartate NADH shuttle (MAS) (23). Song and colleagues reported that the deletion of *A. fumigatus* AgcA caused lowered azole susceptibility (24). They additionally demonstrated that agcA localized in mitochondria (24). As shown in *slc25a12*-deficient mice, MAS activity in mitochondria was impaired significantly, resulting in decreased mitochondrial NADH (25). In the mitochondrial matrix, NADH is the donor of electrons transferred to ubiquinone in complex I (NADH oxidoreductase). As reported by Bowyer (26) and Bromley (27) the disruption of 29.9 kDa subunit in complex I affected azole susceptibility. For that reason, *agcA* disruption might represent a similar effect of subunit disruption by NADH decrease in mitochondria, although further investigation is necessary.

In *A. fumigatus*, UbcD has not been characterized, although the homologs have been studied in *S. cerevisiae* (Ubc4p), and Ubc4 in *Schizosaccharomyces pombe* (Ubc4), which is a ubiquitin-conjugating enzyme E2 (28). In *S. pombe*, Ubc4 cleaved Sre2, a sterol regulatory element-binding protein (SREBP) (29). Fungal SREBPs are involved in adaptation under hypoxic conditions. The cleavage of SREBP promotes adaptation to the environment such as hypoxia (30). *A. fumigatus* has two homologous SREBPs: SrbA and SrbB. These proteins regulate ergosterol biosynthesis and azole susceptibility, as well as hypoxic adaptation. The SREBP-deficient strains showed decreased expression of some genes in the ergosterol biosynthesis pathway including the *cyp51A* gene (31, 32). A plausible mechanism of the azole tolerance led by T98K mutation is that UbcD constitutively activated the function as a ubiquitin ligase, which in turn activates SREBP proteins. An increased expression level of *cyp51A* is expected if this mechanism is supported. However, the expression of *cyp51A* was not higher in TBCZ34 than in the parental strain (data not shown). Generally, the ubiquitin system broadly affects various pathways in an organism, suggesting that the mutation affects protein degradation and cleavage in multiple pathways. The effect of the mutation at the 98th residue remains unresolved.

A mutation in *abcJ* co-occurred with *ubcD* mutation. From the result found for the MIC value against TBCS34/*ubcD*_wt_ strain (Table 2), the contribution of *abcJ* for azole tolerance might be minor in TBCZ34. Further studies must be conducted to elucidate the relation between AbcJ and azole tolerance.

Results of homology analysis show that *rttA* gene was conserved broadly among *Aspergillus* spp. Homologous proteins of some *Aspergillus* spp. such as *A. terreus*, *A. clavatus*, and *A. nidulans* included the C6 zinc finger domain. However, RttA in *A. fumigatus* impairs the domain (Fig. S5). As shown in a dataset provided by Bowyer on FungiDB (33), the transcript of *rttA* was increased twice to three times under the presence of itraconazole, suggesting that the gene plays an important role in adaptation to azole antifungals.

In summary, by *in vitro* selection with TBCZ, we ultimately obtained four TBCZ-adapted isolates. Furthermore, we demonstrated that mutations in MfsD, AgcA, UbcD, and RttA contribute to azole tolerance. These genes are novel players contributing to azole tolerance, although their roles remain unknown. Further studies are expected to be useful to understand the mechanism of azole tolerance in *A. fumigatus*.

## Materials and Methods

### Strains and media

*A. fumigatus* strains used for this study are shown in Table 1 and Table 3. Potato dextrose agar, *Aspergillus* minimal medium (MM) (34), and MOPS-buffered RPMI1640 (RPMI) media were used for culturing those strains. When using uracil auxotrophs, 10 mM uracil was added to appropriate media.

**Table 3.** *Aspergillus fumigatus* strains used for this study except for isolates during selection rounds.

| **Name** | **Genotype** | **Source** |
| --- | --- | --- |
| **AfS35** | *akuA::loxP* | Fungal Genetics Stock Center (strain No. A1159) |
| **$\Delta mfsD$** | *akuA::loxP, mfsD::hph* | This study. |
| **$\Delta agcA$** | *akuA::loxP, agcA::hph* | This study. |
| ***rttA*<sub>A83T</sub>** | *akuA::loxP, rttA*<sub>A83T</sub>, *hph* | *rttA* was replaced with *rttA*<sub>A83T</sub>. *hph* gene cassette was inserted between Afu7g04735 and *rttA*. |
| ***rttA*<sub>wt</sub>** | *akuA::loxP, hph* | *hph* gene cassette was inserted between Afu7g04735 and *rttA*. |
| **$\Delta pyrG$** | *akuA::loxP, \Delta pyrG* | This study, derived from AfS35. |
| **TBCZ34/*ubcD*<sub>wt</sub>** | *akuA::loxP, \Delta pyrG, hph* | *ubcD*<sub>T98K</sub> was replaced with wild-type *ubcD*. |

### Selective passaging with TBCZ

The selection procedure is presented in Fig. 1A. In the initial round, the parental strain *A. fumigatus* Δ*pyrG* (10^5^ conidia) was inoculated in RPMI with 10 mM uracil (RPMI/Ura) and 1/2 MIC of TBCZ, and cultured at 35 °C for 7–10 days at 130 rpm. After culturing, the recovered fungal cells were spread on three plates containing different concentrations of TBCZ. Isolates were picked up and transferred to potato dextrose agar plates with 10 mM uracil. Spore suspensions of each isolate were prepared and stocked at −80 °C until further use. For the next selection round, isolates were mixed. The MIC of TBCZ against the mixture was found (mixed MIC_TBCZ_ in Fig. 1B). We conducted three selection rounds. The 1/2 MIC of TBCZ were found to be 2 and 8 μg/mL, respectively, in the second and third rounds. The MICs were found at 96 h after inoculation according to the modified broth microdilution method, as described previously (35).

### Genome sequencing and detecting mutated points

*A. fumigatus* strains were cultured in potato dextrose broth with 0.1% yeast extract and 10 mM uracil. After culturing at 35°C for 20 hr, mycelia were recovered on Miracloth (Merck KGaA, Darmstadt, Germany). Kits (*Quick*-DNA Fungal/Bacterial; Zymo Research, Irvine, CA, USA) were used for genome DNA preparation. We performed whole-genome sequencing using next-generation methods, as described earlier (6). In brief, we prepared a fragmented DNA library using a NEBNext Ultra II FS DNA Library Prep Kit for Illumina according to the manufacturer’s instructions (New England Biolabs, Ipswich, MA, USA). Sequencing of a paired-end 2 × 150 bp mode on a HiSeq X system (Illumina, San Diego, CA, USA) was done by BGI Japan (Kobe, Japan). For SNP calling, we mapped the reads to the genome sequence of *A. fumigatus* reference strain Af293 (29,420,142 bp, genome version: s03-m04-r03) (36, 37) using BWA version 0.7.15-r1142 (38). Then we identified SNPs using SAMtools ver. 1.8 (39, 40) and filtered them with >10-fold coverage, >30 mapping quality, and 75% consensus using in-house scripts (41, 42). We annotated the functional effect of SNPs with SnpEff ver. 4.1 l (43).

### Sanger sequencing

Sanger sequencing was performed as described elsewhere (44).

### Gene disruption and replacement

DNA fragment preparation by fusion PCR and transformation by protoplasting were done as described in an earlier report (45). Primers used for this study are shown in Table S1. 5-fluoroorotic acid (5-FOA) was used for the selection of the Δ*pyrG* strain (Fig. S1). A cassette containing a hygromycin resistance gene was used to select hygromycin-resistant transformants (46) (Fig. S2). To revert each mutated position, we replaced each gene fragment from the parental strain containing a hygromycin-resistant gene (Fig. S3 and S4). Those correct replacements and disruptions were confirmed by PCR.

### Statistics

Comparisons of colony diameters were analyzed for statistical significance using one-way analysis of variance (ANOVA) followed by Dunnett’s test for post-hoc analysis.

## Acknowledgments

This study was supported by AMED under Grant Number JP19fm0208024 and JP20jm0110015. We thank Kasumi Kodama for technical assistance. We acknowledge DBCLS TogoTV for providing images used in this manuscript. TT, DH, AW, and HT participated in the study design. TT, KO, and MS conducted the experiments of selective passaging with TBCZ, Sanger sequencing, and gene disruption and replacement. YK and HT conducted genome sequencing and detection of mutated points. TT prepared the draft manuscript. TT, DH, AW, and HT mainly reviewed and edited the manuscript. All authors read and approved the final manuscript. The authors declare that they have no competing interests.

